# GPCR-mediated regulation of glial TNF production

**DOI:** 10.64898/2026.06.12.731854

**Authors:** Zahra Abbasi, Shirin Sadighparvar, Marie Franquin, David Stellwagen

**Affiliations:** Department of Neurology and Neurosurgery, Centre for Research in Neuroscience, Research Institute of the McGill University Health Center, Montréal, Québec, Canada

## Abstract

Neuromodulators generally act through G-protein-coupled receptors, but their effects on glia are not well defined. Here we examine the impact of various G-protein-coupled signaling pathways on glia, using the production of the pro-inflammatory cytokine tumor necrosis factor alpha (TNF) as a measure of activation. TNF is a major component of the innate immune response but is also an important regulator of synaptic function and can be released by both astrocytes and microglia. Using pharmacological and chemogenetic approaches, we characterized the response to activation of the Gi, Gq, and Gs signaling pathways in rat astrocyte and microglia cultures and human induced pluripotent stem cells (hiPSCs) derived astrocytes. Across all tested glia, activation of the Gs pathway results in a stark decrease in TNF expression. Similarly, activation of Gq signaling also results in a reduction in TNF mRNA levels. Conversely, Gi activation in astrocytes and microglia increases TNF levels both *in vitro* and *in vivo*. The impacts of GPCRs on TNF production were not consistent for other pro-inflammatory cytokines. Overall, this work demonstrates that G protein-mediated activation and inhibition in glia should be considered separately from the effects seen in neurons.

## INTRODUCTION

Neuron-glia communication is a crucial aspect of brain function and is necessary for maintaining the overall health and homeostasis of the nervous system. It is now commonly acknowledged that non-neuronal glial cells, such as astrocytes and microglia, are essential to normal brain functioning (Adamczyk, 2023), involved in more than just passively supporting neurons (Auld and Robitaille, 2003) and contribute to synaptic development and synaptic function (Eroglu and Barres, 2010).

One way that neurons communicate with glia is through neuromodulators, which in turn can modify the signals transmitted form glial cells to neurons (Fields and Stevens-Graham, 2002). For example, neuronal release of neuromodulators, such as dopamine or serotonin, which can regulate the glial production of the proinflammatory cytokine tumor necrosis factor alpha (TNF) (Duseja et al., 2014; Lewitus et al., 2016). A component of the innate immune system, TNF also serves a regulatory role in non-pathological brain functions (McCoy and Tansey, 2008). It is present in the central nervous system (CNS) under physiologic conditions, where it plays a vital role in regulating synaptic activity (Heir and Stellwagen, 2020; Santello and Volterra, 2012), and can be released by both astrocytes and microglia.

Both types of glia actively contribute to TNF-mediated modifications of brain circuits and behavior, with TNF released by astrocytes mediating homeostatic synaptic plasticity triggered by chronic activity deprivation (Heir et al., 2024; Heir and Stellwagen, 2020; Stellwagen and Malenka, 2006) and microglia produce TNF in reaction to various stimuli including stress (Kemp et al., 2022) and drugs of abuse (Lewitus et al., 2016). The complex interplay of neurons, glial cells, and TNF highlights the necessity of identifying the mechanisms controlling glial TNF production. It has been established that both astrocytes and microglia possess numerous G-protein coupled receptors (GPCRs) for a diversity of neurotransmitters and neuromodulators (Gu et al., 2021; Kofuji and Araque, 2021; Nimmerjahn and Bergles, 2015; Pocock and Kettenmann, 2007). Thus, neuromodulators signaling via GPCRs represents a major route for neuronal-glial communication and likely are important regulators of glial TNF production.

Here we characterized the impact of various G-protein coupled receptors on TNF production from glia. We specifically activated Gi-, Gq-, and Gs-coupled receptor signaling pathways in both astrocytes and microglia, using a combination of pharmacological and Designer Receptors Exclusively Activated by Designer Drugs (DREADDs)-based approaches. In rat astrocyte and microglia cultures and human induced pluripotent stem cells (hiPSCs) derived astrocytes, our findings reveal a consistent pattern of response, with Gs and Gq pathways strongly suppressing glial TNF production. Conversely, Gi activation in astrocytes and microglia increases TNF levels *in vitro* and *in vivo*. These responses were specific for TNF and not other cytokines, and distinct from the typical neuronal response to GPCRs.

## RESULTS

### Norepinephrine decreases TNF production in astrocytes

Astrocytes are responsive to neuronal signals, which impact numerous astrocyte functions including the production of cytokines. As astrocytes are non-excitable cells, we wanted to characterize the glial responses to neuromodulators, acting through G-protein coupled receptors (GPCRs). We previously demonstrated that serotonin signaling can increase TNF mRNA in astrocytes (Duseja et al., 2015), but there are multiple serotonin receptors linked to a variety of G-protein cascades. Here we chose to test the effect of norepinephrine (NE) on the TNF production in astrocytes (Fig 1A). We cultured cortical glia from neonatal rat pups, and used shaking to remove the microglia (McCarthy and De Vellis, 1980), resulting in highly enriched astrocyte cultures (qPCR Ct values: GFAP = 16.8, Iba1 = 34.6). The resultant astrocyte cultures were treated with 50 µM NE for 4 hours, and TNF mRNA analyzed by qPCR (Fig 1B). This treatment markedly reduced TNF mRNA. This is likely due to a decrease in TNF mRNA production as we didn’t observe any changes in the TNF mRNA degradation rate following NE treatment (Fig 1C). Once new RNA synthesis was blocked by actinomycin (5 µg/mL), TNF mRNA was rapidly degraded and was virtually undetectable by 30 minutes; however, this rate was not altered by prior NE treatment. Thus, NE appears to impact TNF mRNA synthesis rather than degradation.

**Figure 1.**
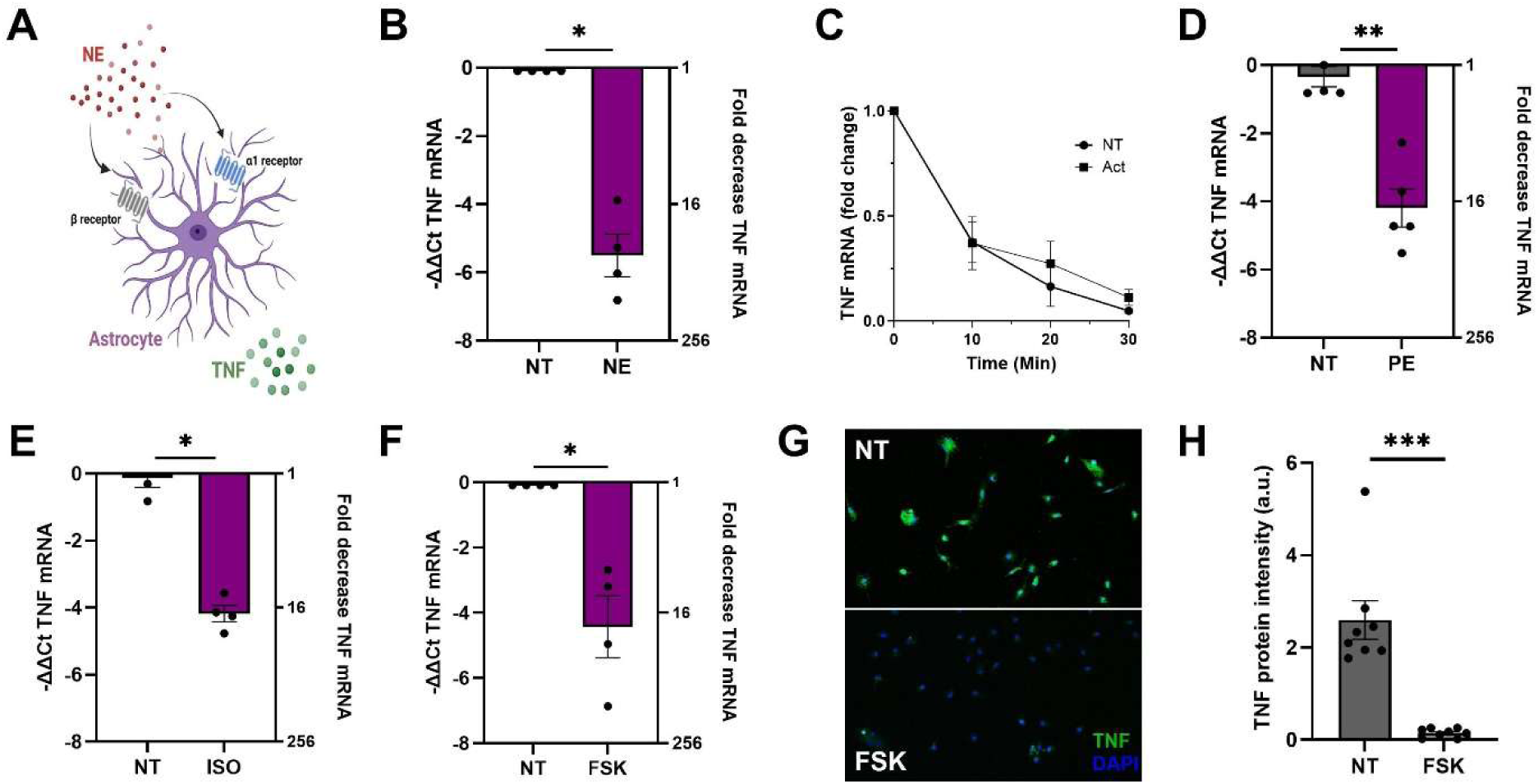
Norepinephrine acting through α- and β-adrenergic receptors reduces TNF mRNA levels in astrocytes. **(A)** Schematic representation of the experimental design, with Norepinephrine (NE) binding to β-adrenergic and α1- adrenergic receptors on astrocytes to influence TNF production. **(B)** Astrocyte cultures treated with 50μM NE for 4h had significantly decreased TNF mRNA levels as measured by qPCR (Mann-Whitney, P=0.0286, n=4). The -ΔΔC_t_ values are plotted on the left axis, while fold change is plotted on the right axis. **(C)** NE treatment did not increase the rate of TNF mRNA degradation. Astrocytes were treated with NE, followed by the RNA synthesis inhibitor actinomycin (5 µg/mL). NE treatment did not alter the rate of TNF RNA degradation (2-way repeated measure ANOVA; effect of treatment: F(1, 16) = 1.708, P = 0.2097; effect of time: F(3, 16) = 34.14, P < 0.0001; interaction: F(3, 16) = 0.734, P = 0.5468). **(D)** The α1-adrenergic receptor agonist Phenylephrine (PE; 50μM, 4h), significantly decreased TNF mRNA levels in astrocytes (Mann-Whitney, P=0.0079, n=5). **(E)** Astrocyte cultures treated with the β-adrenergic receptor agonist Isoprenaline (ISO; 50μM, 4h) also had a similar decrease in TNF mRNA levels (Mann-Whitney, P=0.0286, n=4). **(F)** Astrocyte cultures were treated with the adenylyl cyclase activator Forskolin (FSK; 20μM, 4h) had significantly reduced TNF mRNA levels (Mann-Whitney, P=0.0286, n=4). **(G)** TNF protein was also decreased following 4h FSK treatment. Micrographs of astrocyte cultures immunostained for TNF (green) and DAPI (blue) from non-treated cultures and cultures treated with 20μM FSK. **(H)** Mean intensity was significantly decreased by FSK treatment (Mann-Whitney, P= 0.0002, n=8).

NE can act through α- and β-adrenergic receptors, with both α1- and β-adrenergic receptors expressed on astrocytes (Oe et al., 2020; Perez, 2020). To test the receptors involved, we used Phenylephrine (PE), a selective α1-receptor agonist and Isoprenaline (ISO), a general β-adrenergic receptor agonist. Treating cultured astrocytes with PE (4 hours; 50 µM) also resulted in a significant decrease in TNF mRNA levels (Fig 1D), which was comparable to the effect observed with NE. We then tested the role of β-adrenergic receptors on astrocytes by treating astrocytes with 50 µM ISO for 4 hours (Fig 1E), leading to an equivalent reduction in TNF mRNA levels. These data indicate that the cultured astrocytes used here express both α1- and β-adrenergic receptors.

As all β-adrenergic receptors are Gs-coupled GPCRs that activate adenylyl cyclase to increase cAMP levels, we next verified that increasing cAMP levels would lead to a decrease in TNF mRNA expression. To this end, astrocyte cultures were treated with 20 µM Forskolin (FSK), an adenylyl cyclase activator, for 4 hours. FSK treatment markedly reduced TNF mRNA level in astrocytes (Fig 1F). We have previously observed a strong correlation between TNF mRNA and protein levels (Heir et al., 2024; Lewitus et al., 2016), and we confirmed that the reduction in TNF mRNA following FSK treatment was also reflected in a corresponding decrease in TNF protein levels. Astrocyte cultures were treated with 20 µM FSK for 4 hours, and immunostained for TNF (Fig 1G). We observed a significant reduction in TNF fluorescence signal upon FSK treatment (Fig 1H). Therefore, the increase in cAMP production through Gs-coupled GPCR activation significantly reduces TNF production in astrocytes.

### Gq signaling pathway activation decreases TNF production in astrocytes

Unlike β-adrenergic receptors, α1-receptors are Gq coupled. We have previously shown that glutamate acting through type 1 mGluRs, which are also Gq coupled receptors (Niswender and Conn, 2010), reduces TNF level in astrocyte-only cultures (Heir et al., 2024). To verify that Gq coupled receptor activation reduces TNF expression (Fig 2A), we took advantage of DREAADs, which allow specific activation of a defined G protein coupled pathway (Armbruster et al., 2007). To achieve this, we infected enriched astrocyte cultures with AAV8-gfaABC1D-hM3Gq-mCherry virus, to drive Gq-coupled GPCR expression under a specific astrocyte promoter (Heffernan et al., 2022), and confirmed that a high percentage of the culture was successfully infected and expressing mCherry (Fig 2B). Six days after infection, cultures were treated with the DREADD agonist Clozapine-N-oxide (CNO; 10 µM, 6h) to stimulate Gq activation and TNF mRNA assessed by qPCR (Fig2C). Following CNO treatment, we observed a significant reduction in TNF mRNA level in astrocytes compared to the untreated group. We then confirmed that the decrease in TNF mRNA was accompanied by a decrease in TNF protein. CNO treatment also resulted in a substantial reduction in TNF protein as assessed by immunocytochemistry (Fig 2D-E). This demonstrates that, similarly to Gs activation, activation of the Gq signaling pathway decreases TNF levels.

**Figure 2.**
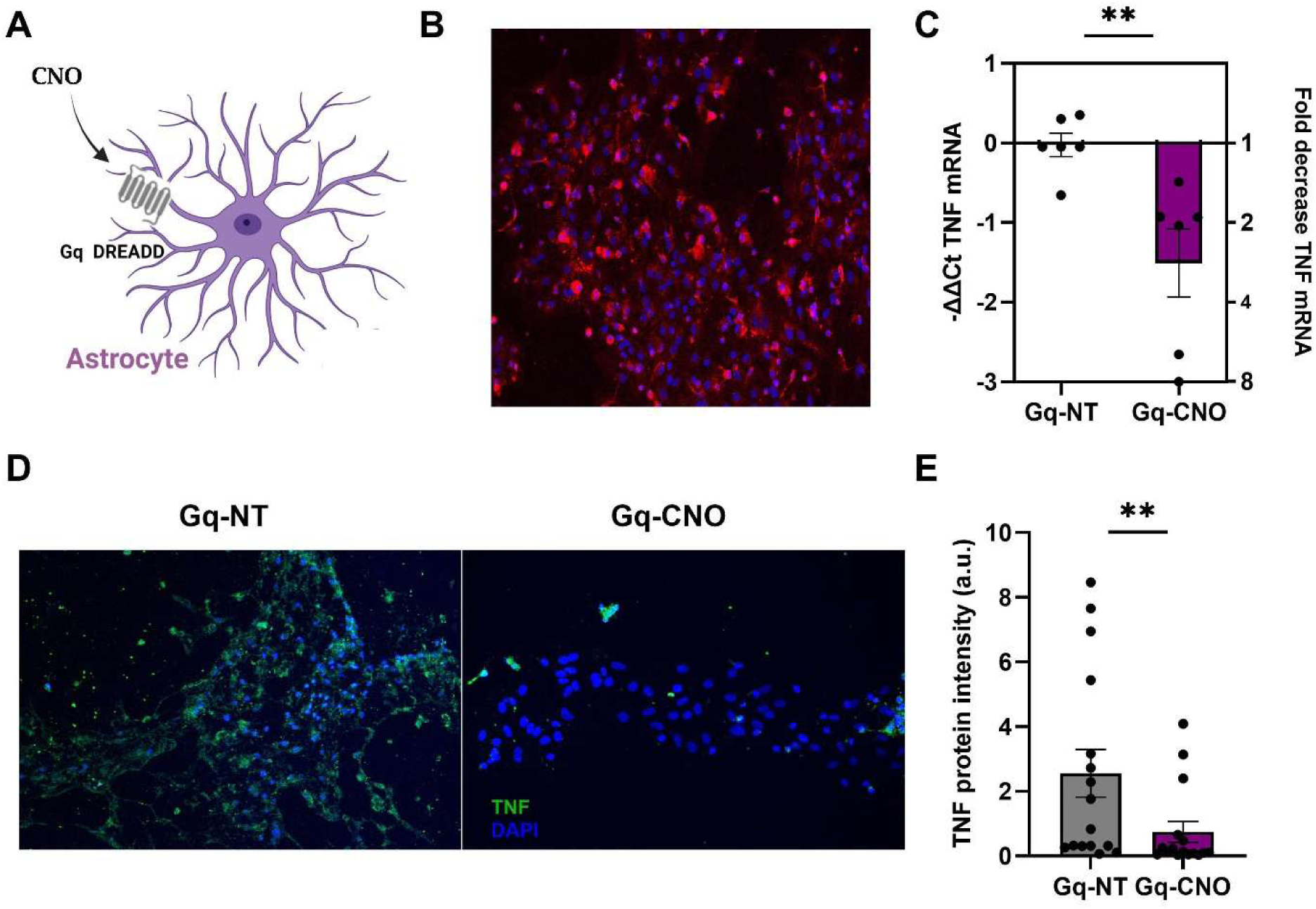
Gq-receptor activation via DREADDs decreased TNF mRNA levels in astrocytes. **(A)** Schematic representation of Gq-DREADD activation in astrocytes using clozapine-N-oxide (CNO) on Gq-DREADD expressing astrocytes, leading to the modulation of TNF production. **(B)** Micrograph showing that the majority of astrocytes were expressing mCherry (red; counterstained with DAPI in blue) six days after being infected with AAV8-gfaABC1D-hM3Gq-mCherry virus. **(C)** Activation of Gq-DREADDS on astrocytes with 10μM CNO for 6h significantly reduced TNF mRNA expression compared with non-treated (NT) DREADD expressing astrocytes (Mann-Whitney, P= 0.0043, n=6). **(D)** Micrographs of TNF protein expression in Gq-DREADD-expressing astrocytes comparing non-treated (NT) and CNO-treated cultures. **(E)** Quantification of TNF protein fluorescent intensity, showing that CNO treatment significantly reduced TNF protein in astrocytes expressing Gq-DREADDs (Mann-Whitney, P=0.0085, n=16).

### Gi signaling pathway activation increases TNF level in astrocytes

We next assessed the effect of the remaining major G-protein cascade, Gi, on TNF production. We have previously demonstrated that dopamine acting through D2 receptors, which are Gi coupled receptors (Beaulieu and Gainetdinov, 2011), increased TNF production in microglia (Lewitus et al., 2016). To determine if Gi signaling caused a similar response in astrocytes (Fig 3A), we expressed a Gi-linked DREADD using AAV2/5-gfaABC1D-hM4D(Gi)-mCherry virus in astrocyte cultures and verified that a high percentage of astrocytes were successfully infected (Fig 3B). As isolated astrocytes already express elevated TNF levels, likely due to lack of neuronal activity, we pre-treated the astrocytes cultures with glutamate (40 µM; 24 hours prior) to mimic neuronal activity and lower TNF levels (Heir et al., 2024) and avoid any ceiling effects. The Gi-DREADD was then activated with 10 µM CNO and TNF mRNA assessed 6h later (Fig 3C). Activating Gi-DREADDs with CNO resulted in a significant increase in TNF levels above non-CNO treated controls. To confirm that the change in mRNA is reflected in a change in protein, we assessed TNF protein by immunocytochemistry in CNO-treated and CNO-untreated cultures (Fig 3D-E), and observed a significant increase in TNF protein with Gi-DREADD activation. Thus, the Gi signaling pathway increases TNF production, unlike the Gs and Gq pathways, in rat astrocytes

**Figure 3.**
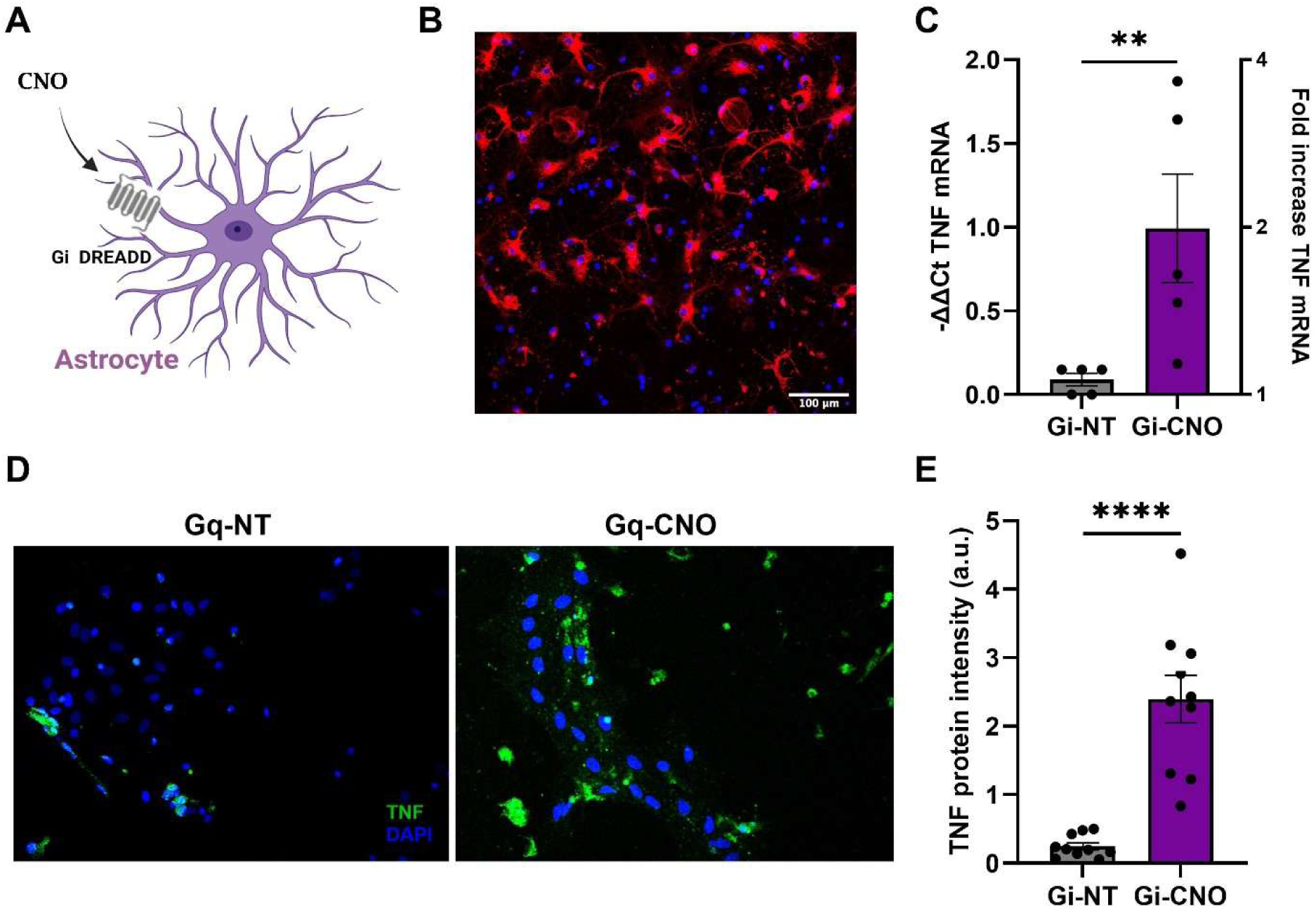
Gi receptor activation using DREADDs increases TNF mRNA levels in astrocytes. **(A)** Schematic representation of Gi-DREADD activation in astrocyte using CNO to activate Gi-DREADDs expressed on astrocytes. **(B)** Micrograph of astrocyte cultures infected with AAV2/5-gfaABC1D-hM4D(Gi)-mCherry virus. Six days post-infection, the majority of astrocytes expressed mCherry (red; counterstained with DAPI in blue). **(C)** Gi-DREADD receptor activation by CNO significantly increased TNF mRNA expression, compared with non-treated DREADD expressing astrocytes (Mann-Whitney, P=0.0079, n=5). Cultures were first treated with 100μM Glutamate for 24h, before being treated with 10μM CNO for 6h. **(D)** Micrographs of TNF protein expression in Gi DREADD-expressing astrocytes comparing non-treated (NT) and CNO-treated cultures. **(E)** Quantification of TNF protein fluorescent intensity, showing that CNO treatment significantly increased TNF protein in astrocytes expressing Gi-DREADDs (Mann-Whitney, P<0.0001, n=10).

### Human astrocytes show the same TNF production pattern as rat astrocytes

Several studies have demonstrated significant differences between human astrocytes and rodent astrocytes in terms of function, gene expression, and morphology (Zhang et al., 2016). Human astrocytes are more numerous, complex, and have unique subclasses compared to rodents (Oberheim et al., 2009), with distinct gene expression responses to stimuli (Hamby et al., 2012) and higher baseline TNF production (Han et al., 2013). Therefore, we tested if TNF production in human astrocytes is regulated similarly to rodent astrocytes.

We differentiated 3 lines of human induced pluripotent stem cells (hiPSCs) into astrocytes using an established protocol (Ponroy Bally et al., 2020). The resultant human astrocytes were positive for mRNA of typical astrocytic markers, S100β and ALDH1L1 (-ΔΔCt S100β = 2.2123 ± 0.9104. T-test p=0.8344, N=4 and -ΔΔCt ALDH1L1 = 1.4333 ± 0.2976. T-test p=0.8825, N=4; data not shown), and had protein expression of GFAP, S100β, and ALDH1L1, and no expression of iPSC and NPC markers such as SOX2 (Fig 4A). To test the G-protein response of these human astrocytes, we first treated the astrocytes with 20μM FSK for 4 hours to investigate the Gs pathway (Fig 4B). This treatment significantly reduced TNF production across all three lines, similar to the effect observed in rat astrocytes. Next, we infected human astrocytes with Gq-DREADDs with the same virus used before. After activating these receptors with 10 μM CNO for 6h, we observed a significant reduction in TNF levels across all individuals (Fig 4C). Finally, to investigate the Gi pathway, we infected the human astrocyte with the Gi-DREADD. As done for testing Gi activation in rat astrocytes, the cultures were pre-treated with 100 µM glutamate for 24h to lower basal TNF levels, prior to CNO application (10 μM; 6h) to activate the Gi-DREADDs. This activation led to a significant increase in TNF levels (Fig 4D). Therefore, GPCR activation led to a similar pattern of TNF production in human astrocytes as in rat astrocytes, although the effects were reduced in amplitude compared with rodents.

**Figure 4.**
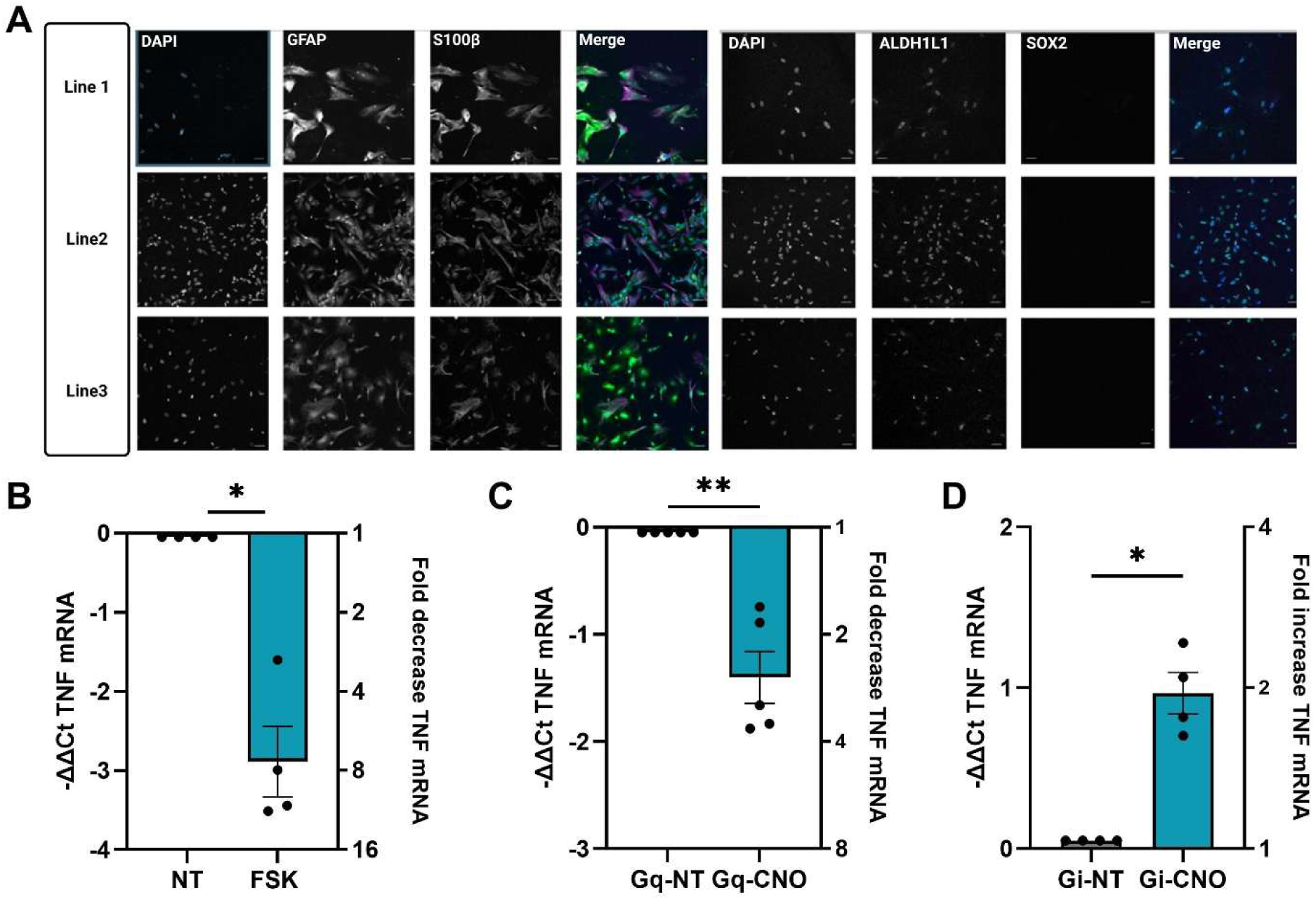
Human iPSC-derived astrocytes have similar responses to GPCRs as rat astrocytes. **(A)** Micrographs showing immunocytochemistry demonstrating that the iPSC-derived astrocytes were positive for the astrocytic markers GFAP, S100β and ALDH1L1 but negative for the NPC marker SOX2. Scale bar (50μm). **(B)** Group data showing that expression of TNF mRNA in human astrocytes decreased following forskolin treatment to activate the Gs pathway (FSK; 20μM, 4h), compared with non-treated sister cultures (Mann-Whitney, P=0.0286, n=4). **(C)** Group data showing that activation of Gq-DREADDs by CNO (10μM, 6h) reduced the expression of TNF mRNA in human astrocytes compared with non-treated sister cultures (Mann-Whitney, P=0.0286, n=5). **(D)** Group data showing that activation of Gi-DREADDs by CNO (10μM, 6h) increased the expression of TNF mRNA in human astrocytes compared with non-treated sister cultures (Mann-Whitney, P=0.0286, n=4).

### Microglia demonstrate a TNF production pattern identical to that of astrocytes

After defining the G-protein regulation of TNF production in astrocytes, we wanted to determine if there was similar regulation in microglia. Microglia are another crucial type of glial cell in the brain, can produce substantial amounts of TNF and play a significant role in neuroinflammation and neurodegenerative diseases (Hanisch, 2002; Smith et al., 2012). To make microglia cultures, we prepared cortical mixed glial cultures from neonatal rat pups and utilized a shaking method to isolate the microglia (McCarthy and De Vellis, 1980), resulted in enriched microglia cultures. As microglia are resistant to viral infection, which often results in low infection efficiency (Maes et al., 2019), we employed a pharmacological strategy here (Fig 5A). To assess the effect of Gs pathway activation on TNF production in microglia, we began with NE treatment. Microglia were exposed to 50μM NE for 4 hours, followed by qPCR to measure TNF mRNA levels. As with astrocytes, microglia demonstrated a significant reduction in TNF levels following NE treatment (Fig 5B). Also similar to astrocytes, the β-adrenergic receptor agonist isoprenaline (ISO; 4h, 50μM) dramatically reduced in TNF mRNA levels in astrocytes (Fig 5C), implicating the Gs pathway. To verify this, we next treated the microglia with the cyclic AMP activator FSK. A 4-hour treatment with 20μM FSK resulted in a significant decrease in TNF production from microglia, mirroring the pattern observed in astrocytes (Fig 5D). To investigate the effect of the Gq signaling pathway on TNF production in microglia, we targeted group I mGluRs, which are Gq-coupled receptors (Niswender and Conn, 2010). We treated microglia with 15 µM DHPG, an agonist for group I mGluRs, for 4h. DHPG treatment significantly reduced the levels of TNF from microglia culture (Fig 5E). To study the Gi pathway in microglia, we chose to target GABA_B_ receptors, which are Gi-coupled receptors (Enna, 2001). We treated microglial cultures with the GABA_B_ receptor agonist baclofen (50 µM, 4h) and observed significantly increased TNF production in microglia, similar to the effects observed in astrocytes (Fig 5F). These data support that G-protein coupled receptor signaling pathways in microglia display a similar pattern to those in astrocytes regarding the regulation of TNF production.

**Figure 5.**
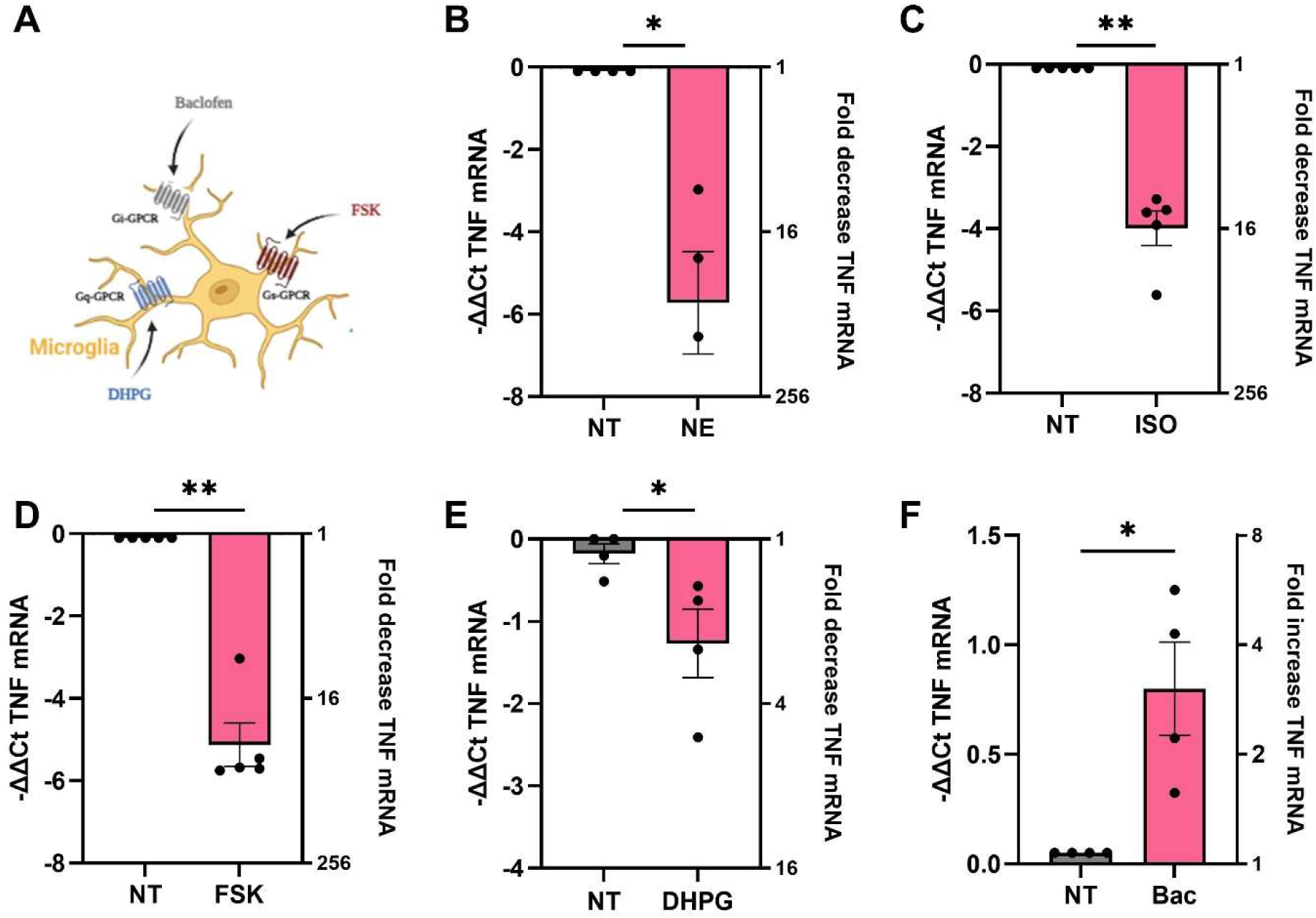
Microglia have an identical pattern of GPCR-regulation of TNF production as astrocytes. **(A)** Schematic representation of the pathways for GPCR activation in microglia by various ligands to determine the impact on TNF mRNA expression. **(B)** Treating enriched microglia cultures with 50μM NE for 4h significantly decreased TNF mRNA levels (Mann-Whitney, P=0.0286, n=4). **(C)** Treating microglia cultures with the β adrenergic receptor agonist Isoprenaline (ISO; 50μM, 4h) also significantly reduced TNF mRNA levels (Mann-Whitney, P=0.0079, n=5). **(D)** Microglia cultures treated with adenylyl cyclase activator Forskolin (FSK; 20μM, 4h) had a similar reduction in TNF mRNA levels (Mann-Whitney, P=0.0079, n=5). **(E)** To test the Gq pathway, microglia cultures were treated with 15 µM DHPG, an agonist of group I mGluRs, for 4h. DHPG treatment significantly decreased TNF mRNA levels in microglia (Mann-Whitney, P=0.0286, n=4). **(F)** To test the Gi pathway, microglia cultures were treated with 50 µM Baclofen, a GABA_B_ receptor agonist, for 4h. Baclofen treatment significantly reduced TNF levels in microglia (Mann-Whitney, P=0.0286, n=4).

### TNF regulation by GPCRs is independent of IL1 regulation

We next wanted to determine if the GPCR regulation of TNF represented a general activation of glia, resulting in the co-regulation of other pro-inflammatory cytokines. Activated microglia typically increase production of a suite of pro-inflammatory cytokines (Hanisch and Kettenmann, 2007), and we have recently shown that brain expression of many immune signaling molecules is regulated by neuronal activity (Heir et al., 2025).

To investigate whether the GPCR-mediated regulation of TNF production extends to other pro-inflammatory cytokines, we examined the expression of interleukin-1α (IL-1α) and interleukin-1β (IL-1β) using the same material and treatments used for TNF expression. Forskolin treatment (to activate the Gs pathway) increased both IL-1α and IL-1β expression in astrocytes (Suppl. Fig 1A,D), which is the opposite of the TNF response. The response to the other pathways in astrocytes via Gq- and Gi-DREADD activation did not lead to significant changes in either IL-1α or IL-1β, and had no clear correlation with the changes observed in TNF expression (Suppl. Fig 1B,E; Suppl. Fig 2C,F). We had similar dissociation of the inflammatory response with microglia. Unlike in astrocytes, forskolin led to a significant decrease in IL-1α (Suppl. Fig2A), although there was a non-significant trend for an increase in IL-1β (Suppl. Fig2D). However, microglia were similar to astrocytes in that there were no significant changes in IL-1α and IL-1β with Gq activation (via DHPG; Suppl. Fig3B,E) or with Gi activation (via baclofen; Suppl. Fig2C,F). Finally, we investigated whether GPCR activation induces general activation in microglia by measuring Iba1 mRNA expression. Activation of GPCRs on microglia with DHPG, baclofen, or forskolin all led to increased Iba1 levels; however, the increase observed with Gs pathway activation via forskolin was not statistically significant (Suppl. Fig 2G-I). These findings suggest that TNF is regulated by distinct mechanisms in response to GPCR signaling in glial, and is not part of a generalized pro-inflammatory response.

### Gi-DREADDs increase astrocyte TNF production in vivo

These results have been from cultured glia, which can have differences from what is seen *in vivo*. Therefore, to verify that GPCR-mediated regulation of glial TNF production occurs *in vivo* and not just *in vitro*, we infected the visual cortex of adult mice with virally-expressed DREADDs. As on-going neuronal activity suppresses astrocyte TNF production (Heir et al., 2024), we chose to use Gi-DREADDs to elevate TNF production rather than Gq-DREADDs to reduce it. Using astrocyte-trageted AAVs, we expressed Gi-DREADDs in one hemisphere of the visual cortex while expressing a control AAV in the countralateral hemisphere. After 3-4 weeks recovery, animals were given the DREADD agonist deschloroclozapine (DCZ; 3 μg/kg; 3h; (Nagai et al., 2020)) and we compared the TNF mRNA levels between the two visual cortices. Activation of Gi-DREADDs in astrocytes resulted in over a 4-fold increase in TNF mRNA relative to the control hemisphere (Fig 6). Thus, Gi activation will increase astrocyte TNF mRNA expression *in vivo* as it does *in vitro*.

**Figure 6.**
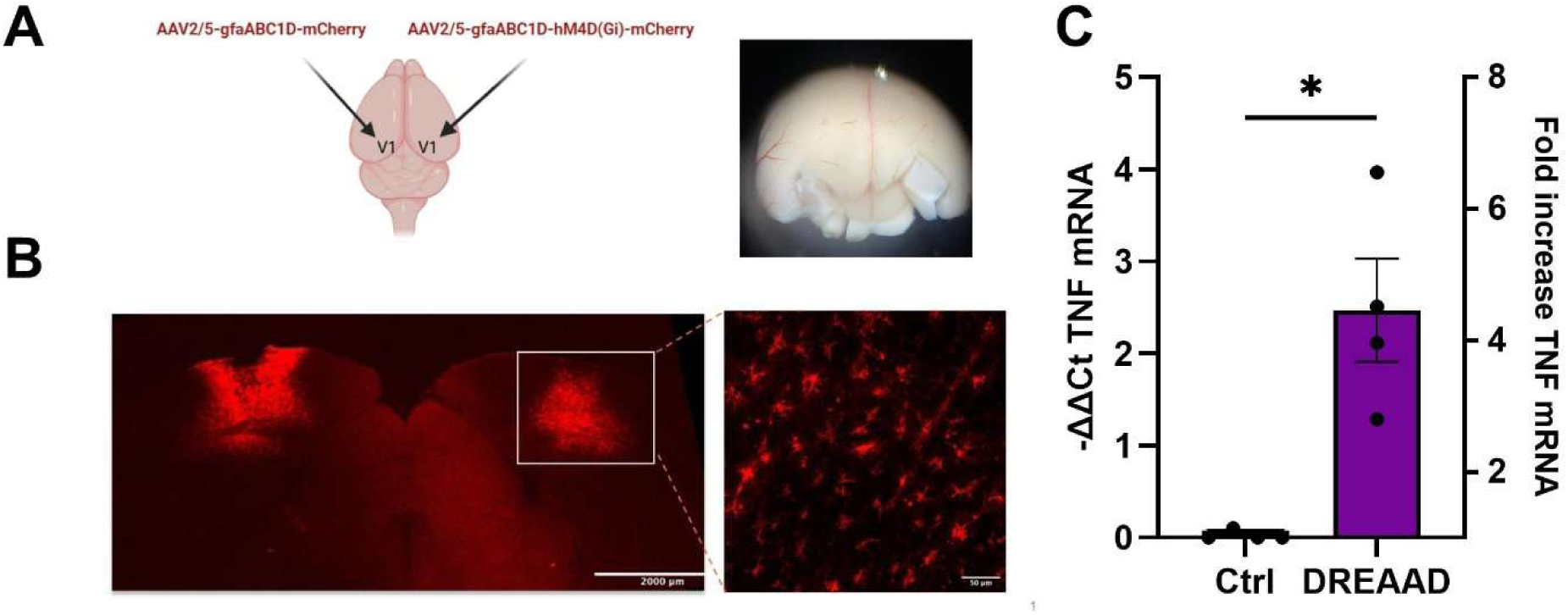
Astrocytic Gi-DREADD activation increases TNF mRNA *in vivo*. (A) Schematic of the injection and treatment protocol for *in vivo* expression of Gi-DREADDs in V1 of one hemisphere and a control virus in the contralateral hemisphere. The micrograph shows the areas dissected for analysis. (B) Micrographs showing the expression of mCherry bilaterally in V1, with a higher magnification showing expression in astrocytes. (C) Following DCZ treatment (3 µg/kg, IP, 3 h), the Gi-DREADD expressing V1 had significantly more TNF mRNA expression than the control V1 (Mann-Whitney, P=0.0286, n=4).

## DISCUSSION

Our study reveals pathway-specific regulation of TNF production by G-protein-coupled receptor (GPCR) signaling in glial cells, identifying a consistent pattern across astrocytes, microglia, and human iPSC-derived astrocytes. Activation of Gs- and Gq-coupled GPCRs suppresses TNF expression, while activation of Gi-coupled receptors increases it (Fig 7). This response is conserved across species and glial subtypes, and was specific for TNF, highlighting a fundamental mechanism by which neuromodulatory signals can shape glial regulatory output to control synaptic modulation in a manner distinct from the GPCR effects commonly seen in neurons.

**Figure 7.**
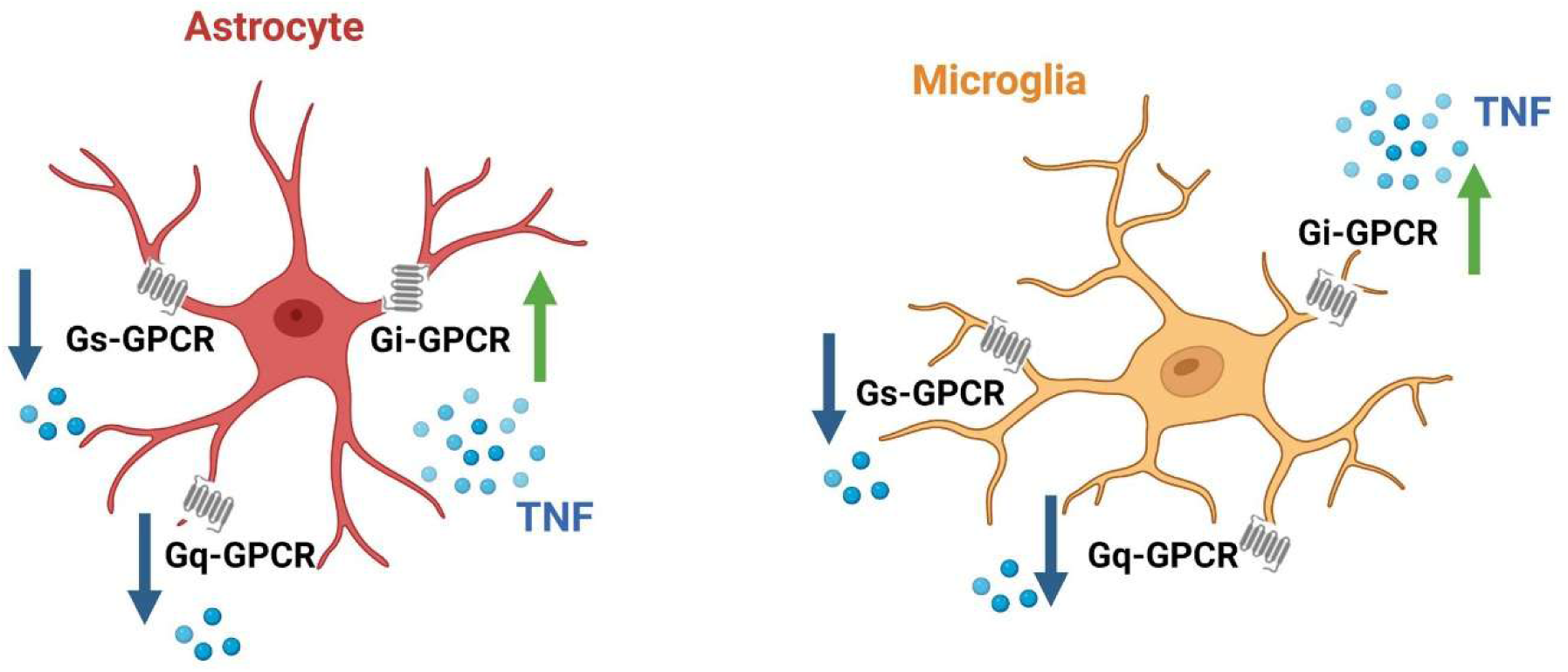
Summary of GPCR activation on TNF expression in astrocytes and microglia. Activation of Gs and Gq pathways in both astrocytes and microglia results in a decrease in TNF levels, while activation of Gi pathways increases TNF levels. This response is the opposite of what occurs in neurons, where Gq and Gs activation typically leads to neuronal activation, and Gi activation results in inhibition.

The responses observed in TNF levels upon GPCR activation indicate that Gs, Gq, and Gi pathways all regulate TNF mRNA. The Gs pathway has been shown to elevate cAMP levels in astrocytes, which in turn influence both their shape and function. Increased intracellular cAMP alters astrocyte morphology (Vardjan et al., 2014) and glucose uptake and glycogenolysis, which in turn contribute to gliotransmission and plasticity (Zhou et al., 2019; Zhou et al., 2021). Interestingly, elevated cAMP also is also reported to reduce astrocytic proinflammatory profile by inhibiting cytokine production (Christiansen et al., 2011; Zhou et al., 2019). These findings align with our results, where Gs pathway activation significantly reduced TNF production in glial cells. This suppression is consistent with the known ability of cAMP/PKA signaling to inhibit NF-κB transcriptional activity (Ye, 2001), providing a direct route by which Gs activation can dampen TNF expression.

Conversely, the Gi pathway inhibits cAMP production and would be expected to give the opposite response as Gs activation. And indeed, selective activation of the Gi pathway led to a robust increase in TNF expression. This is consistent with previous work associating Gi signaling with increased proinflammatory cytokine release (Ye, 2001), and our prior work showing that activation of dopamine D2 receptors on microglia results in elevated levels of TNF (Lewitus et al., 2016). However, Gi-DREADD activation in hippocampal astrocytes in an LPS-induced neuroinflammation mouse model inhibited neuroinflammation by reducing proinflammatory cytokine levels (Kim et al., 2021). These divergent effects suggest that the functional role of Gi signaling in glia may be context-dependent, varying by activation states, as we used unstimulated glia for our studies. Additionally, Gi activation in astrocytes can suppress synaptic output in the dorsolateral striatum (Adermark et al., 2022) and enhanced contextual memory (Nam et al., 2019), but whether these effects are TNF-dependent is unknown. This divergence underscores the complexity of glial GPCR signaling, which may integrate multiple extracellular cues and internal states to determine cytokine output and functional consequences.

Gq activation has similar results as Gs activation, suppressing TNF production although via a distinct mechanism. This is consistent with our previous finding that Gq-coupled group I mGluRs on astrocytes suppress TNF expression (Heir et al., 2024). This contrasts with leukocytes where Gq signaling can promote inflammatory gene expression (Ye, 2001). Gq activation and subsequent Ca²⁺ release also has other effects on glial; Gq-DREADD activation of astrocytes led to a ∼50% increase in evoked miniature excitatory post-synaptic currents (mEPSCs) (Adamsky et al., 2018) and enhanced slow inward currents (SICs) in nearby neurons (Durkee et al., 2019). Moreover, astrocytic Gq-DREADDs are shown to facilitate NMDA-dependent LTP in CA3-CA1 synapses and improve memory processes (Adamsky et al., 2018). These findings highlight how Gq activation in astrocytes and subsequent Ca²⁺ release affect neuronal activity. Long-term activation of this pathway may allow astrocytes to decode neuronal activity to down-regulate TNF, thereby maintaining homeostasis during normal levels of synaptic activity.

To test if GPCR-mediated effects represent general activation of glia, we tested if the regulation of TNF extended to other cytokines by measuring IL-1α and IL-1β expression. Our data indicates that TNF and the IL-1s are regulated differently in astrocytes in response to GPCR activation. For IL-1β, this is consistent with previous reports suggesting that IL-1β and TNF are independently controlled by specific pathways (Dinarello, 2009; Hehlgans and Pfeffer, 2005). While TNF production is often regulated through NF-κB-dependent pathways (Heir et al., 2024), IL-1β production requires caspase-1 activation, highlighting their differential regulation (Brough and Rothwell, 2007). However, IL-1α differs from both TNF and IL-1β in its regulation. It does not require caspase-1 processing and can be released in precursor form, often linked to cell stress or membrane damage (Rider et al., 2011; Werman et al., 2004). The lack of GPCR-mediated effects on IL-1α in our study indicates that it is regulated by pathways distinct from those controlling the other cytokines, consistent with its function as a localized alarmin released in response to cell injury, rather than as a broadly acting neuromodulator. Together, these results indicate that GPCR-mediated modulation of glial cytokines is selective, with certain pathways targeting TNF while leaving other inflammatory mediators unaffected or inversely regulated.

Astrocytes, the most abundant glial cells in the CNS, show significant differences between humans and rodents in function, gene expression, and morphology (Zhang et al., 2016). Human astrocytes are more complex and abundant, with unique subclasses absent in rodents (Oberheim et al., 2009). Studies indicate that rodent astrocytes exhibit distinct gene expression profiles when activated by various stimuli, suggesting species-specific glial responses (Hamby et al., 2012; Zamanian et al., 2012). Despite differences in gene expression profiles between human and rodent astrocytes, we observed a similar GPCR response pattern, at least in terms of TNF production. This conservation suggests that the underlying GPCR–TNF signaling modules are deeply embedded in glial biology, even if other transcriptional outputs vary across species.

To further dissect the mechanism underlying the GPCR-induced reduction in TNF mRNA, we examined whether this effect reflected decreased transcription or enhanced mRNA degradation. Using actinomycin D to block transcription, we confirmed that TNF mRNA in astrocytes is intrinsically short-lived, with the majority degraded within 30 minutes, consistent with earlier reports (Kratochvill et al., 2011; Mijatovic et al., 2000). NE treatment produced did not significantly accelerate decay, suggesting that the suppression of TNF mRNA by NE is driven primarily by reduced transcription, although an effect on transcript stability cannot be excluded.

The present study provides important information on the GPCR signaling pathways regulating TNF production from glia. These findings are not restricted to cultured or immature glia, as *in vivo* activation of DREADDs in adult astrocytes gives similar results. This has significant implications for understanding neuromodulation within the neuron-glia network. Astrocytic TNF is known to mediate homeostatic synaptic plasticity (Heir et al., 2024; Stellwagen and Malenka, 2006), with type I mGluRs contributing to the downregulation of TNF in the presence of adequate neuronal activity (Heir et al., 2024). Other neuromodulators, including serotonin and dopamine (Duseja et al., 2015; Lewitus et al., 2016) can influence neuronal excitability and plasticity through the upregulation or downregulation of glial TNF. This may be an important mechanism for the long-term modulation of neuronal circuits by neuromodulators, and allow neuromodulators to gate the induction of homeostatic plasticity, including how homeostatic plasticity is influenced by sleep-wake states and neuromodulatory tone (Bottorff et al., 2024).

In addition to offering insights into the fundamental roles of GPCRs in neural circuits and their maintenance, a deeper understanding of these pathways may also reveal how these mechanisms are disrupted in pathological conditions. Given that TNF production is a hallmark of many diseases and injury states, understanding the biological processes that may be altered in these situations is crucial. From a translational perspective, selective targeting of these GPCR pathways could offer a means to fine-tune glial TNF in diseases where its dysregulation contributes to pathology, such as neurodegeneration, psychiatric disorders, or chronic pain.

Overall, this study defines a core principle of glial biology: that neuromodulatory inputs, via distinct GPCR pathways, selectively tune TNF expression in astrocytes and microglia, which in turn impacts circuit function. These findings bridge neuroimmune signaling with classic GPCR pharmacology and provide a framework for understanding how glial cells interpret the neuromodulatory landscape to influence brain function and plasticity.

## MATERIALS AND METHODS

### Animals

Sprague Dawley rat pups were acquired from Charles River Laboratories. C57BL/6J mice (JAX000664) were acquired from Jackson laboratories and bred in house. All animal procedures were done according to guidelines of the Canadian Council for Animal Care and overseen by the Montreal General Hospital Facility Animal Care Committee.

### Glial cultures

Glial cultures were made using P0/P1 Sprague Dawley rat pups. The cortex was dissected in cold HBSS media, then treated with 0.05% Trypsin-EDTA (Gibco) for 20 minutes. The tissue was triturated using P1000 pipette tips to make a cell suspension and then plated in T75 cell culture flasks (Sarstedt) with DMEM media (Gibco), supplemented with 10% FBS. To separate astrocytes and microglia, confluent mixed glial cultures were put on an orbital shaker at 37C for 4 hours. The supernatant containing microglia was then passed through 100µm cell strainer and transferred to PDL-coated plates. The cells that remain attached after collecting microglia were astrocytes.

### Generation and maintenance of iPSC lines

Fibroblasts used in this study for reprogramming were from healthy individuals and obtained from the Coriell Institute. The specific lines were as follows:

Line 1: GM19688, 57-year-old Caucasian man

Line 2: ND29194, 51-year-old Caucasian woman

Line 3: GM00321, 40-year-old Caucasian woman

Fibroblasts were reprogrammed using the human episomal iPSC reprogramming kit (ALSTEM, RF202_1) containing the four Yamanaka factors (Takahashi and Yamanaka, 2006) and a puromycin selection cassette. The episomal vector was electroporated using the Neon Transfection System (Invitrogen). Cells were then plated on tissue culture plates coated with Matrigel (Corning) in DMEM plus 10% FBS overnight. Subsequently, the media containing 2 μg/mL puromycin (Sigma-Aldrich) was introduced. 48 hours post-transfection, the medium was changed to TesR-E7 media (Stem Cells Technologies). Upon the formation of iPSC colonies, manual detachment was done using a blunted needle, and the floating colonies were plated on Matrigel-coated plates in mTesR1 media (Stem Cell Technologies) supplemented with 10 μM ROCK inhibitor y-27632 (Sigma-Aldrich). The quality of hiPSCs was assessed with four pluripotency markers (Bell et al., 2019): OCT4, TRA-1-60, SOX2, and stage-specific embryonic antigen-4 (SSEA4). We primarily observed large, round colonies, indicating a lack of differentiation, though some less rounded colonies were also present (Suppl. Fig 3). We also verified the ability of our hiPSC lines to form 3D embryoid bodies (EBs), a key characteristic of pluripotent stem cells (data not shown).

### Conversion of hiPSCs into neural progenitor cells (NPC)

We differentiated iPSCs into NPCs using dual SMAD inhibition. On day 0 the cells were fed with neural induction medium 1 (NIM1) containing: DMEM/F12, N2 (Gibco), B27 (Gibco), BSA (1 mg/mL), SB431542 (10 μM) and noggin (200 ng/mL). On day 8, NIM1 was exchanged for NIM2 containing: DMEM/F12, N2, B27. NIM2 medium was refreshed every day for 5 days. On day 13, Cells were cultured in suspension for 2 to 3 days until they formed visible 3D cell aggregates. These aggregates were then selectively plated onto Matrigel-coated cell culture treated plates in NPC medium. These cells were positive for the NPC markers SOX1, SOX2, PAX6, and Nestin (Suppl. Fig 3).

### Differentiation of NPCs into astrocytes

Folllowing a published protocol (Ponroy Bally et al., 2020), we then differentiated NPCs into astrocytes using small molecules (FGF2, EGF, and CNTF). Once NPCs reached 40-50% confluency, the astrocyte medium 1 (AM1) was added to the cells. This medium contains: DMEM/F-12, FBS (10%), epidermal growth factor (EGF) (20 ng/mL; Genscript) and fibroblast growth factor-2 (FGF2) (20 ng/mL; Cell Signaling). Media was changed three times a week for two weeks. At week 3, the astrocyte medium 2 (AM2) was added to the plates. It contains DMEM/F12, FBS (10%), EGF (20 ng/mL) and FGF2 (20 ng/mL) and ciliary neurotrophic factor (CNTF) (5 ng/mL, Genscript) for an additional 6 weeks. At the beginning of week 9, AM2 was replaced with final differentiation medium (FD) containing: DMEM/F12, FBS (10%) and CNTF (5 ng/mL). FD was switched 3 times a week and cells were passaged as needed for a minimum of 2 weeks.

### Drug treatments

L-glutamic acid (100 µM; Tocris) was solubilized in water and added to astrocyte cultures for 24h. Norepinephrine (50 µM; Tocris), Isoprenaline hydrochloride (50 µM; Sigma), and Phenylephrine hydrochloride (50 µM; Tocris) were dissolved in water and added to the astrocyte cultures for 4h. Forskolin (20 µM; Sigma) was solubilized in ethanol was added to astrocytes for 4h, with appropriate vehicle controls. Clozapine N-oxide (CNO) at a final concentration of 10 µM was applied to cultures for 6h following DREADDs expression. (RS)-3,5-DHPG (20 µM; Tocris) and baclofen (50 µM, Tocris) were solubilized in water and added to microglia culture for 4h.

### Viral infection with DREADDs

AAV2/5-gfaABC1D-hM4D(Gi)-mCherry and AAV8-gfaABC1D-hM3(Gq)-mCherry viruses (Neurophotonics, Universite Laval) were directly applied to the astrocyte cultures. A total of 2.75 × 10¹¹ GC was used for each 10 cm culture dish. The culture medium was changed 24h after the virus treatment. mCherry expression was confirmed by immunofluorescent staining 6 days post infection.

### qPCR

Cells were collected from the plates using 0.05% Trypsin-EDTA (Gibco) and mRNA was extracted from the cells by the RNeasy Mini extraction kit (Qiagen). Subsequently, mRNA was transcribed into cDNA using 1 µg input utilizing the Quantitect reverse transcription kit (Qiagen). qPCR was done using the Fast SYBR green master mix (Applied Biosystems) and the StepOne Plus instrument (Applied Biosystems). Primer sequences and annealing temperatures were:

rat TNF: 60.5°C annealing, 600 nM

fwd- CTT CTG TCT ACT GAA CTT CGG G

rev- CTA CGG GCT TGT CAC TCG

rat IL-1β: 54.8 °C annealing, 342 nM

fwd- TGC AGG CTT CGA GAT GAA C

rev- GGG ATT TTG TCG TTG CTT GTC

rat IL-1α: 56.2 °C annealing

fwd- CGC TTG AGT CGG CAA AGA AAT

rev- AGA CAG ATG GTC AAT GGC AGA

rat IBA1: 59 °C annealing

fwd- GCC TCA TCG TCA TCT CCC Ca

rev- AGG AAG TGC TTG ATC CCA

rat GAPDH: 61.5 °C annealing, 350 nM

fwd- TTG TGG ATC TGA CAT GCC G

rev- TGG GAG TTG CTG TTG AAG TC

Human TNF: 54.9 °C annealing

fwd- ACT TTG GAG TGA TCG GCC

rev- CTC AGC TTG AGG GTT TGC TA

Human GAPDH: 54.9 °C annealing

fwd- ACA TCG CTC AGA CAC CAT G

rev- TGT AGT TGA GGT CAA TGA AGG G

All samples were run as triplicates and normalized with GAPDH as a housekeeping gene. All quantification was done using the ΔΔC_t_ method.

### Immunofluorescent staining

For TNF immunostaining, protein export was blocked for 4h with Protein Transport Inhibitor Cocktail (eBioscience), to prevent the cleavage and release of TNF protein. Cells were fixed with 4% paraformaldehyde for 20 min, followed by three times washing with PBS-0.05% tween. Cells were then incubated with blocking buffer (2% NGS, 2% BSA, 0.1% triton in PBS) for 1h, then primary antibody overnight at 4 degrees, followed by appropriate secondary antibody for 1h at room temperature and DAPI labeling (to label cell nuclei).

Primary antibodies used in this study are listed as follows:

Anti-TNF antibody (1:100, Abcam, AB215188), anti-GFAP antibody (1:500, Synaptic Systems, 173 004), anti-S100β antibody (1:500, Sigma, AMAB91038), anti-ALDH1L1 antibody (1:200, Neuromab, 75-164).

All the characterizations for hiPSCs after reprogramming were performed using PSC 4-marker ICC kit (Oct4, Sox2, SSEA4, TRA-1-60; 1:50, ThermoFisher, A24881). All the characterizations for NPCs were done using NPC 4-marker ICC kit (Sox1, Sox2, Nestin, PAX6; 1:50, ThermoFisher, A24354).

### Image Acquisition and Analysis

All images were acquired at 20X on an Olympus FV1000 confocal microscope using identical settings for all images. Images were then transferred to ImageJ for subsequent analysis. For analysis of the TNF labeling, the mean fluorescence intensity (MFI) was quantified for individual images, and then normalized for cell density using DAPI labeling.

### In vivo viral injections

Adult wildtype mice underwent bilateral stereotaxic injections at two sites per hemisphere to achieve viral expression across both superficial and deeper cortical layers of V1. The first injection targeted deeper cortical layers (approximately layers IV–V) at AP −3.5 mm, ML ±2.5 mm, DV −0.5 to −0.7 mm relative to bregma. The second injection targeted a more superficial and slightly rostral region near the V1 boundary at AP −3.3 mm, ML ±2.2 mm, DV −0.3 to −0.5 mm. A control virus (AAV2/5-GFAP-mCherry) was injected into the left hemisphere, whereas AAV2/5-gfaABC1D-hM4Di-mCherry (Neurophotonics, Universite Laval) was injected into the right hemisphere. For each injection, 100nL was infused at 10 nL/min, and the injection needle was left in place for 5–10 min after each injection to reduce backflow along the needle tract. Mice were allowed to recover for 3–4 weeks to permit viral expression. They then received deschloroclozapine (DCZ; 3 μg/kg, i.p.) 3 h prior to V1 dissection, snap freezing, and qPCR processing. TNF mRNA of the DREADD-injected V1 was normalized to that of the contralateral V1 infected with the control virus.

### Statistics

All the results are reported as the mean ± SEM. TNF fluorescent quantification and qPCR data were non-parametric and analyzed by Mann-Whitney tests. Statistical analyses of qPCR data are conducted on -ΔΔC_t_ values, as ΔΔC_t_ values provides a linear output of the assay but fold change is considered more straightforward to comprehend and was also used for the graphs.

## Conflict of Interest statement

The authors declare no competing financial interests

## Acknowledgements

We thank Hooman Salahi and Alex Trottier for technical assistance. This work was supported by the Canadian Institutes for Health Research (CIHR), the Natural Sciences and Engineering Research Council of Canada (NSERC), and the Fonds de Research du Quebec Sante (FRQS).

## SUPPLEMENTAL FIGURES

**Supplementary Figure 1.**
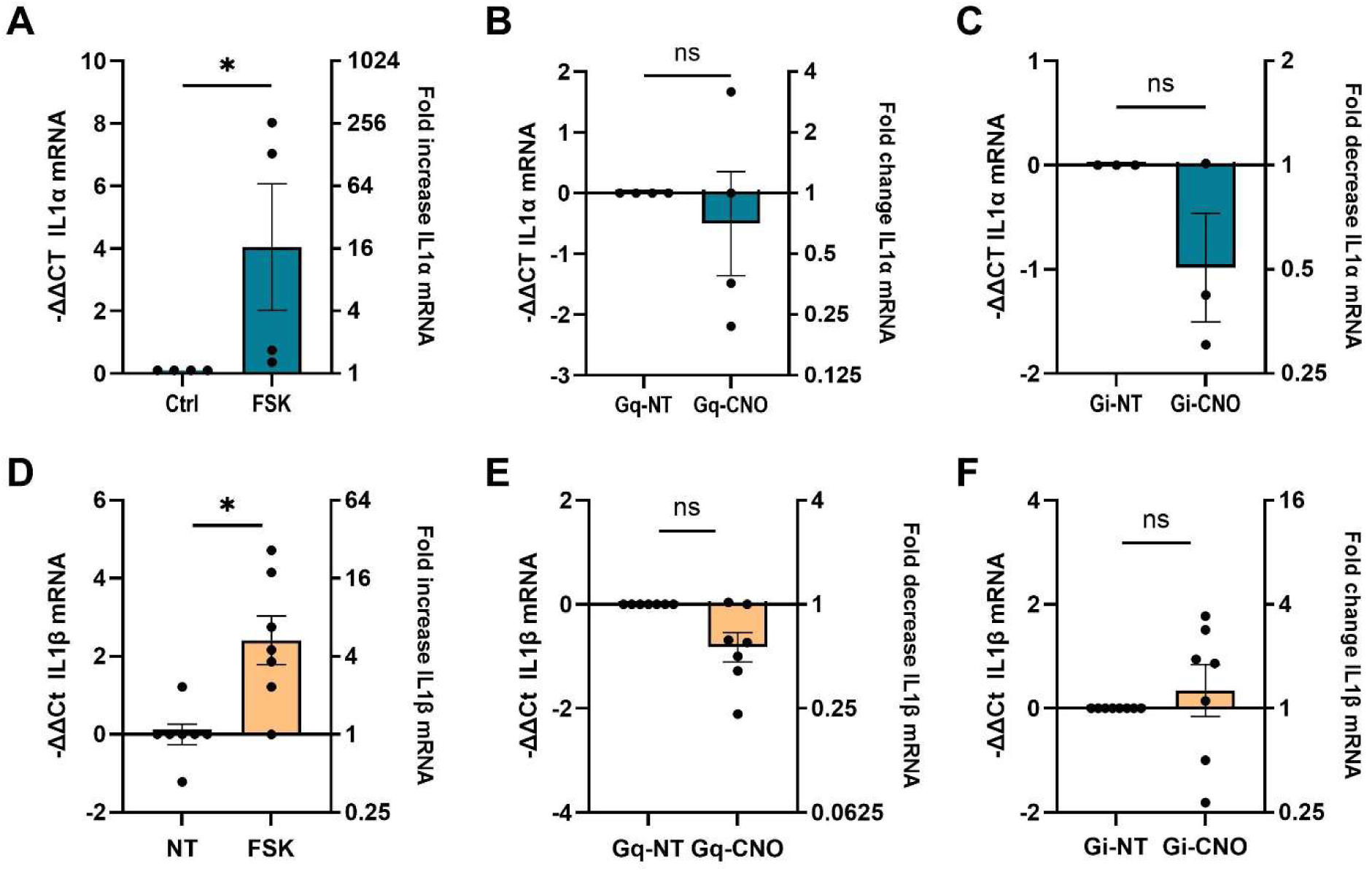
Other pro-inflammatory cytokines were not co-regulated with TNF by GPCRs in astrocytes. **(A)** IL-1α mRNA expression was increased by forskolin treatment (FSK; 20μM, 4h) in cultured astrocyte cultures (Mann–Whitney, P=0.0286, n=4). **(B)** Activation of Gq-DREADD receptors on astrocytes with CNO treatment (10 μM, 6h) did not significantly change IL-1α mRNA (Mann–Whitney, P>0.9999, n=4). **(C)** Activation of Gi-DREADD receptors on astrocytes with CNO did not lead to significant changes in IL-1α mRNA (Mann–Whitney, P=0.7000, n=3). **(D)** FSK treatment (20 μM; 4h) increased IL1β mRNA levels in astrocytes (Mann–Whitney, P=0.0227, n=7), as it did with IL-1α. **(E)** Activation of Gq-DREADD receptors in astrocytes with CNO did not lead to significant changes in IL-1β (Mann–Whitney, P=0.1801, n=7). **(F)** Activation of Gi-DREADD receptors in astrocytes with CNO did not lead to significant changes in IL-1β (Mann–Whitney, P=0.1462, n=7).

**Supplementary Figure 2.**
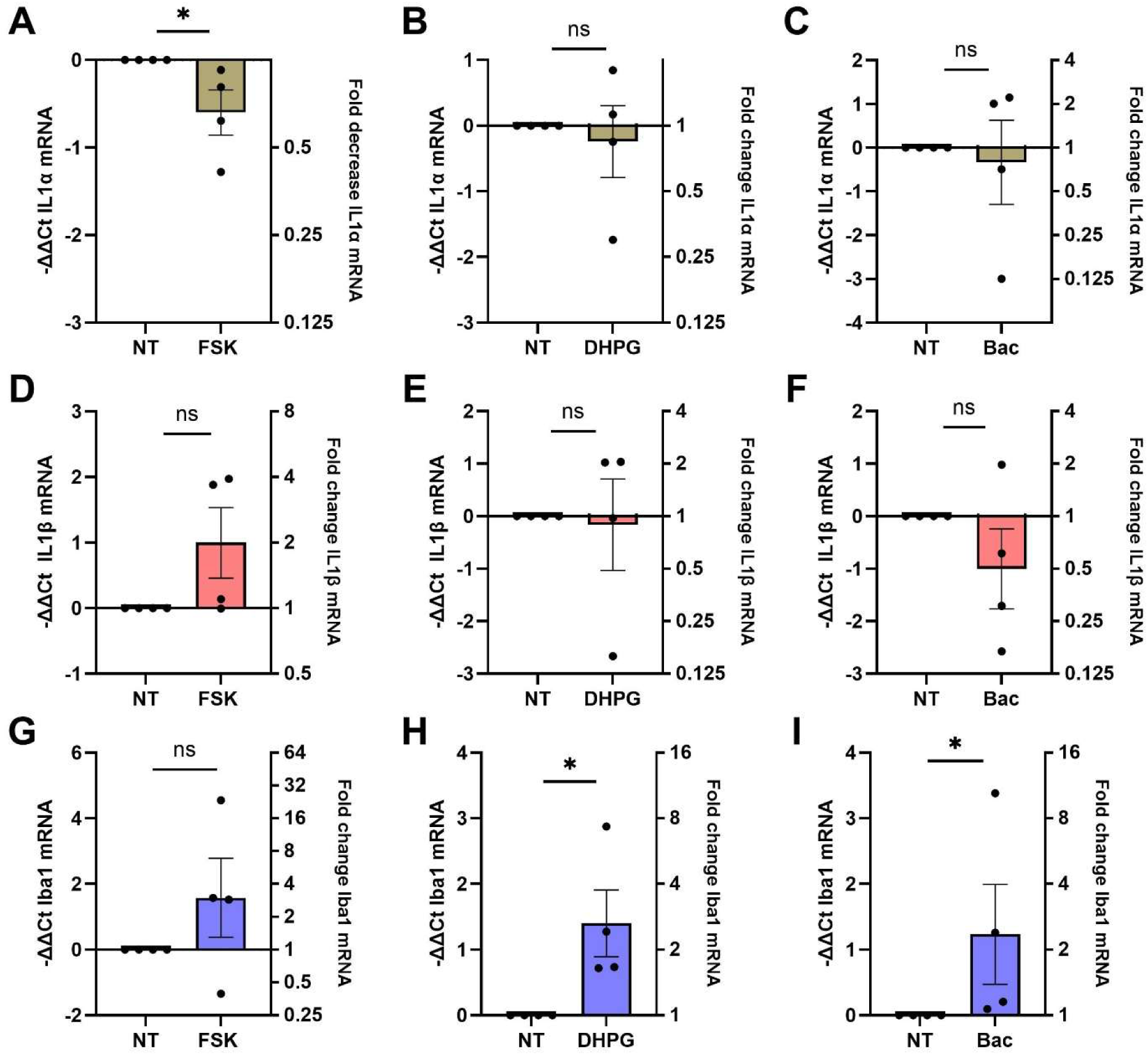
Assessment of other cytokines and activation markers in microglia. **(A)** Microglia cultures were treated with 20μM Forskolin (FSK) for 4h, followed by qPCR to measure IL1a mRNA level. Forskolin led to a significant decrease in IL-1α (Mann–Whitney, P=0.0286, n=4) **(B)** Microglia cultures were treated with 15μM DHPG for 4h, followed by qPCR to measure IL1a mRNA level. No specific pattern of response was observed following Gq activation with DHPG (Mann–Whitney, P>0.9999 n=4). **(C)** Microglia cultures were treated with 50μM Baclofen for 4h, followed by qPCR to measure IL1a mRNA level. No specific pattern was observed following Gi activation with Baclofen (Mann–Whitney, P>0.9999, n=4). **(D)** Microglia cultures were treated with 20μM Forskolin (FSK) for 4h, followed by qPCR to measure IL1b mRNA level. There was a non-significant trend for an increase in IL-1β (Mann–Whitney, P=0.3143, n=4). **(E)** Microglia cultures were treated with 15μM DHPG for 4h, followed by qPCR to measure IL1β mRNA level. No specific pattern was observed following Gq activation with DHPG (Mann–Whitney, P>0.9999, n=4). **(F)** Microglia cultures were treated with 50μM Baclofen for 4h, followed by qPCR to measure IL1β mRNA level. No specific pattern was observed following Gi activation with Baclofen (Mann–Whitney, P=0.3143, n=4). **(G)** Microglia cultures were treated with 20μM Forskolin (FSK) for 4h, followed by qPCR to measure Iba1 mRNA level as a microglia activation marker. Forskolin led to a non-significant increase in Iba1 mRNA level (Mann–Whitney, P=0.3143, n=4). **(H)** Microglia cultures were treated with 15μM DHPG for 4h, followed by qPCR to measure Iba1 mRNA level. DHPG led to a significant increase in Iba1 mRNA level (Mann–Whitney, P=0.0286, n=4). **(I)** Microglia cultures were treated with 50μM Baclofen for 4h, followed by qPCR to measure Iba1 mRNA level. Baclofen also led to a significant increase in Iba1 mRNA level (Mann–Whitney, P=0.0286, n=4).

**Supplementary Figure 3.**
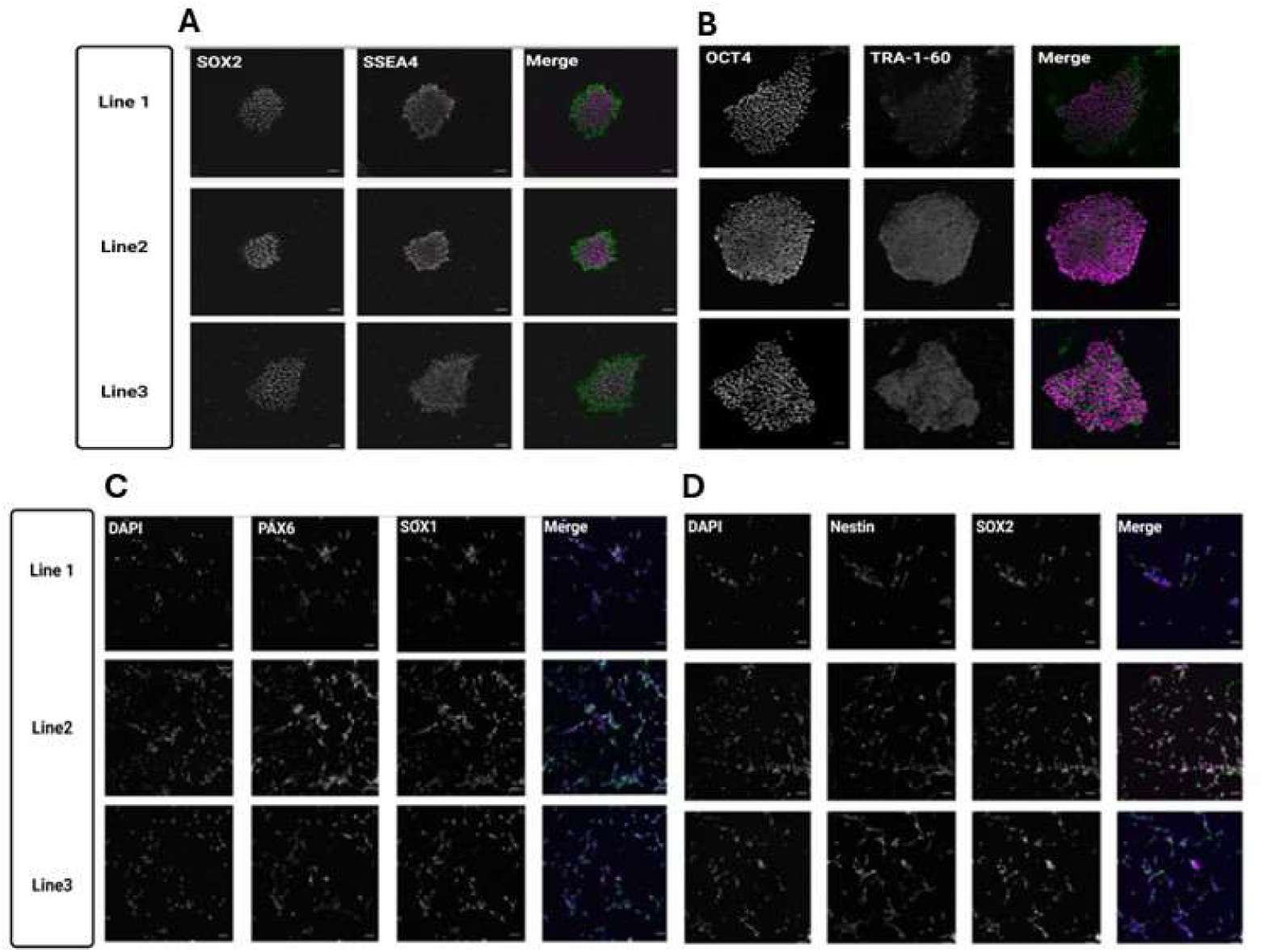
**(A)** iPSC quality control for colonies reprogrammed from fibroblasts. All three iPSC lines exhibited typical round morphology and expressed pluripotency markers including Sox2 and SSEA4. **(B)** IPSCs also expressed pluripotency markers Oct4 and TRA-1-60. **(C)** NPC quality control for neural progenitor cells differentiated from iPSC colonies. All three lines expressed the NPC markers PAX6 and SOX1. **(D)** The NPCs also expressed Nestin and SOX2.

## Notes

### Competing Interest Statement

The authors have declared no competing interest.

## REFERENCES

Adamczyk, A. (2023). Glial-Neuronal Interactions in Neurological Disorders: Molecular Mechanisms and Potential Points for Intervention. Int J Mol Sci 24.

Adamsky, A., Kol, A., Kreisel, T., Doron, A., Ozeri-Engelhard, N., Melcer, T., Refaeli, R., Horn, H., Regev, L., and Groysman, M. (2018). Astrocytic activation generates de novo neuronal potentiation and memory enhancement. Cell 174, 59–71. e14.

Adermark, L., Lagström, O., Loftén, A., Licheri, V., Havenäng, A., Loi, E.A., Stomberg, R., Söderpalm, B., Domi, A., and Ericson, M. (2022). Astrocytes modulate extracellular neurotransmitter levels and excitatory neurotransmission in dorsolateral striatum via dopamine D2 receptor signaling. Neuropsychopharmacology 47, 1493–1502.

Armbruster, B.N., Li, X., Pausch, M.H., Herlitze, S., and Roth, B.L. (2007). Evolving the lock to fit the key to create a family of G protein-coupled receptors potently activated by an inert ligand. Proceedings of the National Academy of Sciences 104, 5163–5168.

Auld, D.S., and Robitaille, R. (2003). Glial cells and neurotransmission: an inclusive view of synaptic function. Neuron 40, 389–400.

Beaulieu, J.-M., and Gainetdinov, R.R. (2011). The physiology, signaling, and pharmacology of dopamine receptors. Pharmacological reviews 63, 182–217.

Bell, S., Hettige, N.C., Silveira, H., Peng, H., Wu, H., Jefri, M., Antonyan, L., Zhang, Y., Zhang, X., and Ernst, C. (2019). Differentiation of human induced pluripotent stem cells (iPSCs) into an effective model of forebrain neural progenitor cells and mature neurons. Bio-protocol 9, e3188–e3188.

Bottorff, J., Padgett, S., and Turrigiano, G.G. (2024). Basal forebrain cholinergic activity is necessary for upward firing rate homeostasis in the rodent visual cortex. Proceedings of the National Academy of Sciences of the United States of America 121, e2317987121.

Brough, D., and Rothwell, N.J. (2007). Caspase-1-dependent processing of pro-interleukin-1β is cytosolic and precedes cell death. Journal of cell science 120, 772–781.

Christiansen, S.H., Selige, J., Dunkern, T., Rassov, A., and Leist, M. (2011). Combined anti-inflammatory effects of β2-adrenergic agonists and PDE4 inhibitors on astrocytes by upregulation of intracellular cAMP. Neurochemistry international 59, 837–846.

Dinarello, C.A. (2009). Immunological and inflammatory functions of the interleukin-1 family. Annual review of immunology 27, 519–550.

Durkee, C.A., Covelo, A., Lines, J., Kofuji, P., Aguilar, J., and Araque, A. (2019). Gi/o protein-coupled receptors inhibit neurons but activate astrocytes and stimulate gliotransmission. Glia 67, 1076–1093.

Duseja, R., Heir, R., Lewitus, G.M., Altimimi, H.F., and Stellwagen, D. (2014). Astrocytic TNFalpha regulates the behavioral response to antidepressants. Brain, behavior, and immunity.

Duseja, R., Heir, R., Lewitus, G.M., Altimimi, H.F., and Stellwagen, D. (2015). Astrocytic TNFα regulates the behavioral response to antidepressants. Brain, behavior, and immunity 44, 187–194.

Enna, S. (2001). GABAB receptor signaling pathways. In Pharmacology of GABA and glycine neurotransmission (Springer), pp. 329–342.

Eroglu, C., and Barres, B.A. (2010). Regulation of synaptic connectivity by glia. Nature 468, 223–231.

Fields, R.D., and Stevens-Graham, B. (2002). New insights into neuron-glia communication. Science 298, 556–562.

Gu, C., Chen, Y., Chen, Y., Liu, C.F., Zhu, Z., and Wang, M. (2021). Role of G Protein-Coupled Receptors in Microglial Activation: Implication in Parkinson’s Disease. Front Aging Neurosci 13, 768156.

Hamby, M.E., Coppola, G., Ao, Y., Geschwind, D.H., Khakh, B.S., and Sofroniew, M.V. (2012). Inflammatory mediators alter the astrocyte transcriptome and calcium signaling elicited by multiple G-protein-coupled receptors. Journal of Neuroscience 32, 14489–14510.

Han, X., Chen, M., Wang, F., Windrem, M., Wang, S., Shanz, S., Xu, Q., Oberheim, N.A., Bekar, L., Betstadt, S., et al. (2013). Forebrain engraftment by human glial progenitor cells enhances synaptic plasticity and learning in adult mice. Cell Stem Cell 12, 342–353.

Hanisch, U.K. (2002). Microglia as a source and target of cytokines. Glia 40, 140–155.

Hanisch, U.K., and Kettenmann, H. (2007). Microglia: active sensor and versatile effector cells in the normal and pathologic brain. Nature neuroscience 10, 1387–1394.

Heffernan, K.S., Rahman, K., Smith, Y., and Galvan, A. (2022). Characterization of the GfaABC1D promoter to selectively target astrocytes in the rhesus macaque brain. Journal of neuroscience methods 372, 109530.

Hehlgans, T., and Pfeffer, K. (2005). The intriguing biology of the tumour necrosis factor/tumour necrosis factor receptor superfamily: players, rules and the games. Immunology 115, 1–20.

Heir, R., Abbasi, Z., Komal, P., Altimimi, H.F., Franquin, M., Moschou, D., Chambon, J., and Stellwagen, D. (2024). Astrocytes are the source of TNF mediating homeostatic synaptic plasticity. The Journal of Neuroscience: the Official Journal of the Society for Neuroscience, e2278222024–e2278222024.

Heir, R., Abuzgaya, M., Adaidi, H., Franquin, M., Abbasi, Z., and Stellwagen, D. (2025). A screen of immune signaling molecules regulated by neuronal activity identifies interferon-gamma as a modulator of synaptic function and anxiety-like behavior. Brain, behavior, and immunity 130, 106066.

Heir, R., and Stellwagen, D. (2020). TNF-Mediated Homeostatic Synaptic Plasticity: From in vitro to in vivo Models. Front Cell Neurosci 14, 565841.

Kemp, G.M., Altimimi, H.F., Nho, Y., Heir, R., Klyczek, A., and Stellwagen, D. (2022). Sustained TNF signaling is required for the synaptic and anxiety-like behavioral response to acute stress. Mol Psychiatry.

Kim, J.H., Rahman, M.H., Lee, W.H., and Suk, K. (2021). Chemogenetic stimulation of the Gi pathway in astrocytes suppresses neuroinflammation. Pharmacology Research & Perspectives 9, e00822.

Kofuji, P., and Araque, A. (2021). G-Protein-Coupled Receptors in Astrocyte-Neuron Communication. Neuroscience 456, 71–84.

Kratochvill, F., Machacek, C., Vogl, C., Ebner, F., Sedlyarov, V., Gruber, A.R., Hartweger, H., Vielnascher, R., Karaghiosoff, M., and Rülicke, T. (2011). Tristetraprolin-driven regulatory circuit controls quality and timing of mRNA decay in inflammation. Molecular systems biology 7, 560.

Lewitus, G.M., Konefal, S.C., Greenhalgh, A.D., Pribiag, H., Augereau, K., and Stellwagen, D. (2016). Microglial TNF-α suppresses cocaine-induced plasticity and behavioral sensitization. Neuron 90, 483–491.

Maes, M.E., Colombo, G., Schulz, R., and Siegert, S. (2019). Targeting microglia with lentivirus and AAV: Recent advances and remaining challenges. Neuroscience letters 707, 134310.

McCarthy, K.D., and De Vellis, J. (1980). Preparation of separate astroglial and oligodendroglial cell cultures from rat cerebral tissue. The Journal of cell biology 85, 890–902.

McCoy, M.K., and Tansey, M.G. (2008). TNF signaling inhibition in the CNS: implications for normal brain function and neurodegenerative disease. J Neuroinflammation 5, 45.

Mijatovic, T., Houzet, L., Defrance, P., Droogmans, L., Huez, G., and Kruys, V. (2000). Tumor necrosis factor-α mRNA remains unstable and hypoadenylated upon stimulation of macrophages by lipopolysaccharides. European Journal of Biochemistry 267, 6004–6012.

Nagai, Y., Miyakawa, N., Takuwa, H., Hori, Y., Oyama, K., Ji, B., Takahashi, M., Huang, X.P., Slocum, S.T., DiBerto, J.F., et al. (2020). Deschloroclozapine, a potent and selective chemogenetic actuator enables rapid neuronal and behavioral modulations in mice and monkeys. Nature neuroscience 23, 1157–1167.

Nam, M.-H., Han, K.-S., Lee, J., Won, W., Koh, W., Bae, J.Y., Woo, J., Kim, J., Kwong, E., and Choi, T.-Y. (2019). Activation of astrocytic μ-opioid receptor causes conditioned place preference. Cell reports 28, 1154–1166. e1155.

Nimmerjahn, A., and Bergles, D.E. (2015). Large-scale recording of astrocyte activity. Curr Opin Neurobiol 32, 95–106.

Niswender, C.M., and Conn, P.J. (2010). Metabotropic glutamate receptors: physiology, pharmacology, and disease. Annual review of pharmacology and toxicology 50, 295–322.

Oberheim, N.A., Takano, T., Han, X., He, W., Lin, J.H., Wang, F., Xu, Q., Wyatt, J.D., Pilcher, W., and Ojemann, J.G. (2009). Uniquely hominid features of adult human astrocytes. Journal of Neuroscience 29, 3276–3287.

Oe, Y., Wang, X., Patriarchi, T., Konno, A., Ozawa, K., Yahagi, K., Hirai, H., Tsuboi, T., Kitaguchi, T., Tian, L., et al. (2020). Distinct temporal integration of noradrenaline signaling by astrocytic second messengers during vigilance. Nat Commun 11, 471.

Perez, D.M. (2020). α1-Adrenergic receptors in neurotransmission, synaptic plasticity, and cognition. Frontiers in pharmacology 11, 581098.

Pocock, J.M., and Kettenmann, H. (2007). Neurotransmitter receptors on microglia. Trends in neurosciences 30, 527–535.

Ponroy Bally, B., Farmer, W.T., Jones, E.V., Jessa, S., Kacerovsky, J.B., Mayran, A., Peng, H., Lefebvre, J.L., Drouin, J., and Hayer, A. (2020). Human iPSC-derived Down syndrome astrocytes display genome-wide perturbations in gene expression, an altered adhesion profile, and increased cellular dynamics. Human molecular genetics 29, 785–802.

Rider, P., Carmi, Y., Guttman, O., Braiman, A., Cohen, I., Voronov, E., White, M.R., Dinarello, C.A., and Apte, R.N. (2011). IL-1α and IL-1β recruit different myeloid cells and promote different stages of sterile inflammation. The Journal of Immunology 187, 4835–4843.

Santello, M., and Volterra, A. (2012). TNFα in synaptic function: switching gears. Trends in neurosciences 35, 638–647.

Smith, J.A., Das, A., Ray, S.K., and Banik, N.L. (2012). Role of pro-inflammatory cytokines released from microglia in neurodegenerative diseases. Brain research bulletin 87, 10–20.

Stellwagen, D., and Malenka, R.C. (2006). Synaptic scaling mediated by glial TNF-α. Nature 440, 1054–1059.

Takahashi, K., and Yamanaka, S. (2006). Induction of pluripotent stem cells from mouse embryonic and adult fibroblast cultures by defined factors. Cell 126, 663–676.

Vardjan, N., Kreft, M., and Zorec, R. (2014). Dynamics of β-adrenergic/cAMP signaling and morphological changes in cultured astrocytes. Glia 62, 566–579.

Werman, A., Werman-Venkert, R., White, R., Lee, J.-K., Werman, B., Krelin, Y., Voronov, E., Dinarello, C.A., and Apte, R.N. (2004). The precursor form of IL-1α is an intracrine proinflammatory activator of transcription. Proceedings of the National Academy of Sciences 101, 2434–2439.

Ye, R.D. (2001). Regulation of nuclear factor κB activation by G-protein-coupled receptors. Journal of Leukocyte Biology 70, 839–848.

Zamanian, J.L., Xu, L., Foo, L.C., Nouri, N., Zhou, L., Giffard, R.G., and Barres, B.A. (2012). Genomic analysis of reactive astrogliosis. Journal of neuroscience 32, 6391–6410.

Zhang, Y., Sloan, S.A., Clarke, L.E., Caneda, C., Plaza, C.A., Blumenthal, P.D., Vogel, H., Steinberg, G.K., Edwards, M.S., and Li, G. (2016). Purification and characterization of progenitor and mature human astrocytes reveals transcriptional and functional differences with mouse. Neuron 89, 37–53.

Zhou, Z., Ikegaya, Y., and Koyama, R. (2019). The astrocytic cAMP pathway in health and disease. International journal of molecular sciences 20, 779.

Zhou, Z., Okamoto, K., Onodera, J., Hiragi, T., Andoh, M., Ikawa, M., Tanaka, K.F., Ikegaya, Y., and Koyama, R. (2021). Astrocytic cAMP modulates memory via synaptic plasticity. Proceedings of the National Academy of Sciences 118, e2016584118.

